# Estimating the time structure of descending activation that generates movements at different speeds

**DOI:** 10.1101/2022.03.21.485078

**Authors:** Rachid Ramadan, Cora Hummert, Jean Stephane Jokeit, Gregor Schöner

**Affiliations:** Institute for Neural Computation, Ruhr University Bochum, Bochum, Germany

## Abstract

In targeted movements of the hand, descending activation patterns must not only generate muscle activation, but must also adjust spinal reflexes from stabilizing the initial to stabilizing the final postural state. We estimate descending activation patterns that change minimally while generating a targeted movement within a given movement time based on a model of the biomechanics, muscle dynamics, and the stretch reflex. The estimated descending activation patterns predict human movement trajectories quite well. Their temporal structure varies across workspace and with movement speed, from monotonic profiles for slow movements to non-monotonic profiles for fast movements. Descending activation patterns at different speeds thus do not result from a not mere rescaling of invariant templates, but reflect varying needs to compensate for interaction torques and muscle dynamics. The virtual attractor trajectories, on which active muscle torques are zero, lie within reachable workspace and are largely invariant movements when represented in end-effector coordinates. Their temporal structure along movement direction changes from linear ramps to “N-shaped” profiles with increasing movement speed.

**Author summary:** What descending patterns of activation drive targeted movements? Based on a model that includes biomechanics, muscles dynamics and the stretch reflex we estimate the descending activation patterns by minimizing their change using optimal control, given a movement target and movement time. The resulting activation patterns predict experimentally observed human movements reasonably well. The temporal structure of the estimated descending activation patterns depends on the speed of the movement, varying from monotonic for slow to non-monotonic for fast movements. This structure reflects the need to compensate for interaction torques and muscle properties. From the model we are able to estimate the virtual attractor trajectories on which all active muscle torques are zero. These lie within reachable workspace, and are relatively uniform across workspace. Their time structure varies from linear ramps for slow movements to N-shaped temporal profiles for fast movements.

## Introduction

Understanding the neural basis of movement generation continues to be a scientific challenge. In fact, movement may be the area of human performance, for which neurophysiological knowledge provides the least constraints [1]. The difficulty derives, in part, from the fact that so many different neural structures contribute substantially to movement generation including multiple cortical areas, the basal ganglia, the cerebellum, and the spinal cord. Moreover, they do so in closed loop, both in the sense of recurrent neural circuits and in the sense of sensory feedback loops. This makes it more difficult to understand the functional properties of neural circuits for motor control than, say, the functional properties of forward neural maps in sensory areas.

The challenges intrinsic to motor neurophysiology are one reason why computational approaches to motor control are so attractive [2–4]. These make it possible to characterize the system of motor control at a “competence” level that abstracts from the underlying mechanisms while still enabling a mapping of such functional descriptions onto different neural structures [5]. This has provided a very productive research framework. Broadly speaking, principles of optimal control and optimal estimation have been used to characterize how the motor control system takes into account learned properties of the the plant and its physical environment. This work has uncovered sophisticated competences that can be described as “internal models” of external contingencies. The adaptation to external force-fields is the lead paradigm for this line of work [6, 7]. Optimal feedback control has been used as a computational framework to characterize how feedback may be used optimally by the motor system to deal with external perturbation and the inherent variability of force production by muscles [8, 9]. This line of thinking has been used to explain patterns of co-variation that may reflect control priorities.

How do we move beyond the characterization of performance achieved in the computational approach to motor control? How may we understand the specific way neural mechanisms shape motor control in light of these computational principles? Here we begin by showing how the descending activation patterns that generate movement are shaped by the stretch reflex. Previous theoretical work has shown that spinal reflexes may change the nature of the control problem [10]. A limitation of such work is that the contribution reflexes make to muscle activation depends on the patterns of descending activation that “command” the movement. How may we identify those patterns at the same time as we assess the contribution of reflexes to motor control? The key idea of our contribution is to use the principles of optimal control to estimate the patterns of descending activation that drive the motor periphery including its reflex loops. We do this in a simple setting, based on a model of the biomechanics of the moving limb, of the dynamics of muscle force generation, and of the stretch reflex [11].

What kind of cost function may provide a plausible constraint to estimate the descending activation pattern? Previous theoretical work has argued that evolutionary optimization would plausibly minimize some measure of “effort” as movement tasks in the world are achieved [12]. Effort was assessed based on a change signal that is non-zero only during movement generation. In its most physiological (and thus non-linear) form, the optimal feedback control problem minimized a change signal directly related to muscle activation, which is zero before the movement, and which returns to zero after the movement [13].

When the stretch reflex is taking into account, the descending “command” is not a change signal because it also contains a component that reflects the shifted postural state achieved at the end of the movement. This is illustrated in Figure 1 for a shoulder extension movement. Prior to the movement, the flexor muscle is activated only to the extent that it opposes and extensor activation, reflecting co-contraction of flexors and extensors. As the extension movement is initiated, flexor may drop. At the end of the movement, it may return to its initial level of activation (if co-contraction is equivalent in the new posture). A descending “command”, which is simply descending activation sent to the motor neurons that drive the flexor muscle, may first drop at the beginning of the extension movement, inducing the muscle’s drop in activation. Because the flexor muscle lengthens during the movement, the afferent signals from flexor muscle spindles increase over the movement. Through the reflex loop, this signal would activate the muscle, resisting the movement. Descending activation must drop and remain at a lowered level at the end of the movement, so that it cancels the increased afferent signal and muscle activation may return to its non-movement state. This is solves the movement-posture problem [14]. Behavioral signatures of a persistent component of the descending activation pattern have recently been reported in the force-field adaptation paradigm [15].

**Fig 1.**
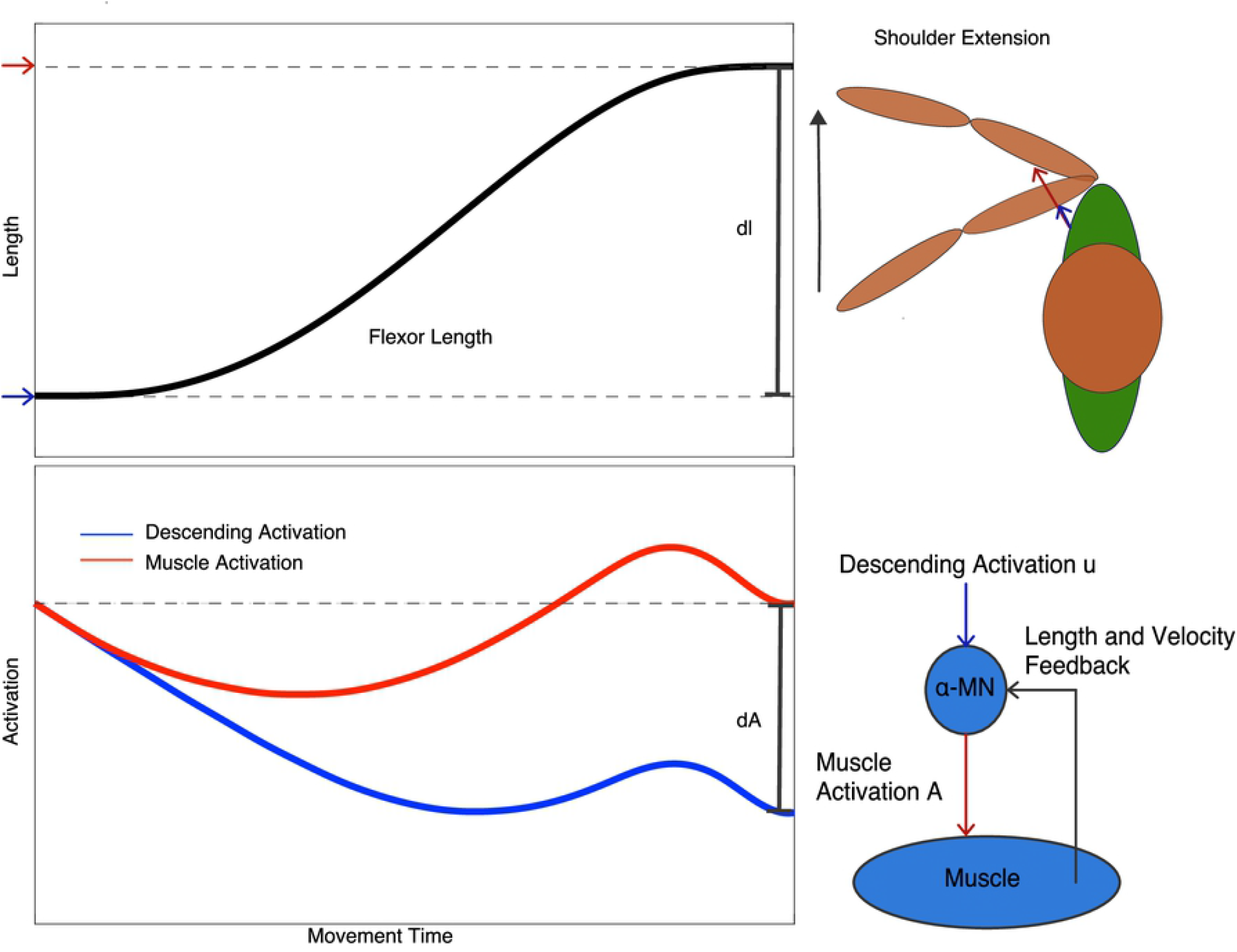
In a shoulder extension movement (illustrated on the top right), the shoulder flexor lengthens (top left). In a simple model of the stretch reflex (illustrated on the bottom right), activation that descends from the brain is combined with length and velocity feedback sensory signals from muscle spindles. When these summed inputs exceed threshold, muscle activation is induced. During shoulder extension, flexor muscle activation first drops from an initial level that reflects the level of co-contraction of flexor and extensor muscles, then rises, contributing to the breaking of the movement, and finally returns to its initial value. Because the flexor lengthens, the feedback signal is larger at the end of the movement than initially. The descending activation pattern is lowered by a matching amount, so that feedback and descending activation cancel, reducing muscle activation and suppressing the flexor’s resistance to the new posture. All curves are from the model, which finds the descending activation pattern that changes minimally to achieve the new posture within a given movement time (see Results Section).

The pattern of descending activation is, therefore, not purely change signal, but includes a tonic component that reflects the new postural state. Descending activation itself is not a good candidate for establishing a cost function for movement generation. We propose instead that the cost function depends on the rate of change of descending activation, but definition a change signal. We evaluate that hypothesis by comparing the kinematics predicted by minimizing this cost function with empirical movement data.

Our main interest lies not in introducing yet another cost function to describe movement competence, however. Our goal is to use this principle primarily to estimate (or rather: provide a good guess of) the descending activation pattern that is consistent with movement of limbs that are endowed with the stretch reflex. This enables us to investigate how descending activation patterns vary across workspace and across different movement speeds. Given the nonlinear nature of interaction torques, varying movement speed changes their relative contribution during movement. We aim, therefore, to directly observe in the temporal patterns of descending activation signatures of internal models that reflect compensation for interaction torques.

This optimal estimate of the temporal patterns of descending activation will also enable us to observe to which extent and over which time interval muscle activation is directly driven by the descending signals. Does the reflex loop contribute to muscle activation during the movement or only once the new postural state has been reached? The stretch reflex defines, in principle, an equilibrium muscle length that determines the postural state at the end of the movement [16]. Does the minimal descending activation pattern define a hypothetical equilibrium configuration of the limb only toward the end of the movement? In the model, we will be able to compute the *attractor trajectory* [17], on which the limb would move entirely passively as established in experiments in which movement was made passive by a finely attuned external force field [17]. Examining the predicted attractor trajectory across workspace and different movement speeds provides insight into the functional significance of the optimal descending activation patterns.

## Materials and methods

### Experimental methods

To obtain experimental data for comparison to the theoretical predictions, we performed a standard movement experiment in the manner of [18]. We aimed at naturalistic movements, restraining movement to planar, two-joint action by instruction rather than by mechanically constraining the arm with a manipulandum.

Eight movements were chosen to sample a large area of the workspace (Figure 2). These movements entail different combinations of shoulder and elbow flexion and extension, with varied roles of the biarticulatory muscles.

**Fig 2.**
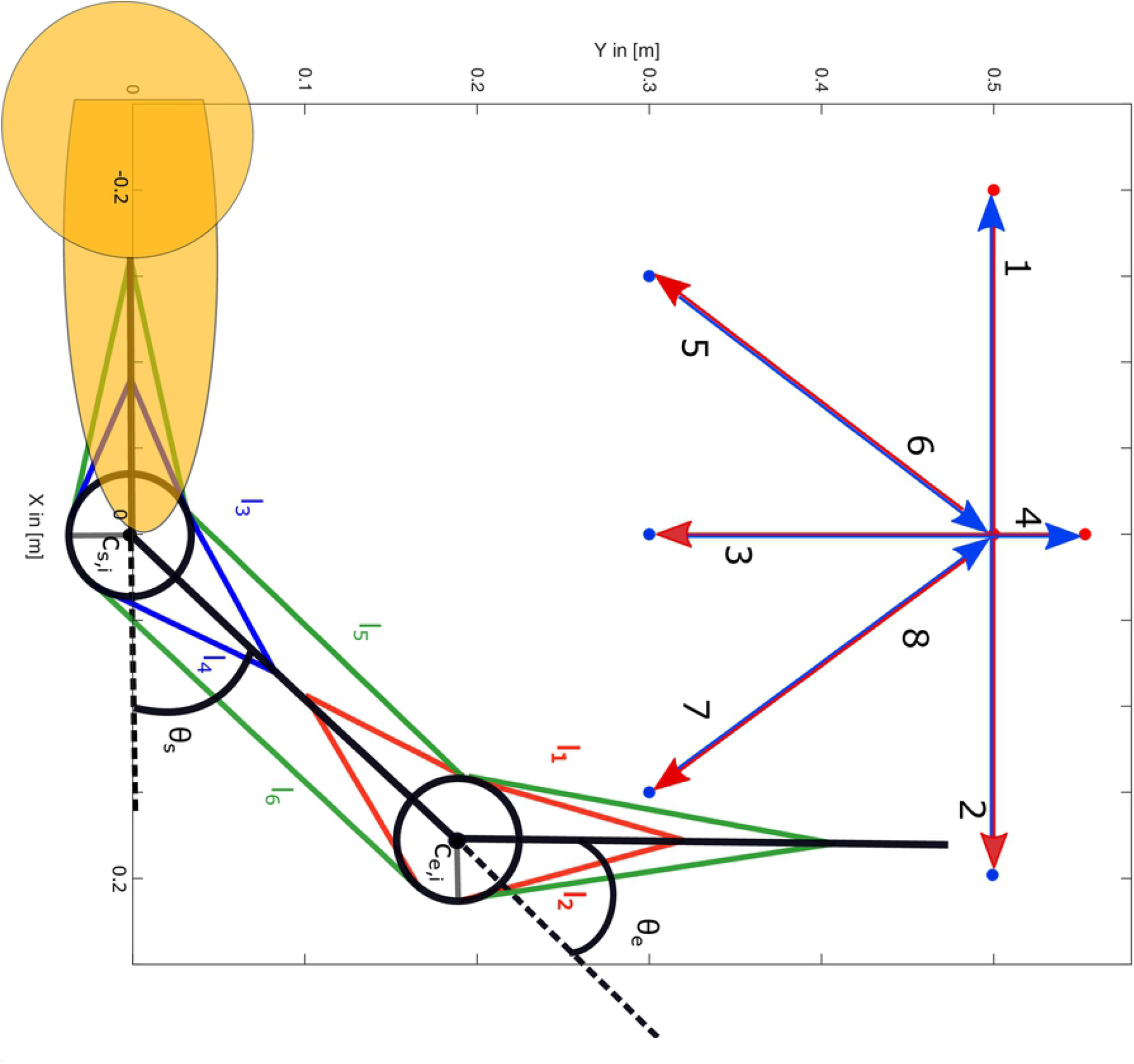
Top: Eight movement conditions (the labels 1-8 are positioned near the end-point of each movement) sample the space of planar movements performed with two degrees of freedom, shoulder and elbow. Bottom: The model includes shoulder and elbow joints in the horizontal plane, articulated by two mono-articular elbow joint muscles (red), two mono-articular shoulder muscles (blue) and two bi-articular muscles that span elbow and shoulder joint (green). The illustrated geometry leaves muscle lever arms invariant across work space.

### Participants

Twelve participants successfully completed the task. (A thirteenth participant was excluded due to failure to complete enough valid trials, see below.) All participants (8 female, 4 male, mean age = 25.67 years, *SD* = 3.80, age range: 22-35) were right handed and had given informed and written consent. The participants were compensated with 10 Euro for the one hour experimental session. The study has been approved by the Ethikkommission am Institut für Neuroinformatik der Ruhr-Universität Bochum.

### Material and procedures

Participants performed point-to-point movements with their right (dominant) hand in the horizontal plane. The movements were captured with the VisualEyez Motion Capture system. Four single chip LED markers were fixated on the arm of the participants, one each at the shoulder and elbow and two on a splint that fixated the wrist (compare figure 2). The two markers at the wrist were attached between the thumb and the index finger at a distance of 2 cm from each other. The right shoulder was strapped to the chair.

### Experimental protocol

The eight examined movements included two movements that covered a distance of 40 cm (movement 1 and 2, comparable to movements used in [11]) and six movements that covered a distance of 25 cm. The movements had varying involvement of the two joints. The experiment started with 10 calibration trials, three to estimate joint angles for different arm positions and seven to calibrate the end-effector position for each target. The experimental trials were organized in two blocks, across which two different levels of movement time were imposed (see below). Within each block, movements were randomized. Each block started with a training trial of each of the eight movements, followed by five sessions of 16 trials, two for each movement, for a total of 10 repetitions per movement and movement time. Overall the participants conducted 176 trials.

During training trials, participants learned to achieve a target movement time by attending to an auditory metronome. The metronome gave two audio signals of different pitch (750 and 550 Hz). The time interval between the two tones signaled the desired movement time, 400 ms for the *fast* condition and 800 ms for the *slow* condition. The two tones were repeated three times, followed by a third tone (620 Hz) that indicated the beginning of the trial. Before the experimental trials, the first two audio signals were played only once before the movement was initiated.

### Data analysis

The motion data was analyzed in *Matlab*. The trajectories were filtered with a third-order Butterworth filter with a cut-off frequency of 5 Hz. Movement onset was defined as the point in time at which end-effector velocity first exceeded 5% of peak velocity and end-effector acceleration exceeded 5% of maximal acceleration. Movement offset was determined as the point in time when end-effector velocity fell below 5% of peak velocity and end-effector deceleration fell below 5% of peak deceleration.

Trials whose movement duration exceeded the median movement duration by more than one third were discarded. If more than four trials for one movement had to be discarded, that movement was deemed unsuccessful and the participant was excluded from the results (and this happened for the thirteenth participant).

The trajectories were time normalized across all experimental trials for each movement and movement time condition. The normalized trajectories were shifted so that the shoulder is at [0,0] and the joint angles were calculated from the shifted trajectories. In calculating joint angle velocities and accelerations, we corrected for time normalization.

Muscle torques, *T*_*j*_, at either joint (*j* = elbow and shoulder) were estimated based on the mean trajectories of each participant using the equation of motion 3 [18]:

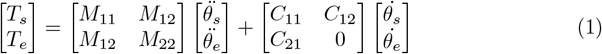

where *M* is the inertial matrix and *C* contains the Coriolis and centripetal torques. Both depend on the joint configuration. Interaction torques, *τ*_*j*_, were obtained as:

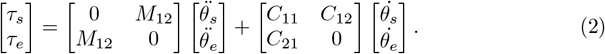

### Theoretical methods

We model the reflex control and force generation of a set of six muscles that actuate the shoulder and elbow joints of a planar arm as illustrated in Figure 2. The model largely follows [11], with a few simplifications outlined below. The model takes six descending, time-varying motor commands *u*_*desc*_(*t*). We determine the “minimal” descending activation pattern by minimizing the amount of change in the descending command needed to transition from the initial to the target posture in a given movement time.

### The neuromusclar model

#### Biomechanics

The same biomechanical equations of motion listed above describe the modelled arm dynamics in terms of the joint angle vector, 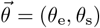 (elbow, e, and shoulder, s) and its derivatives:

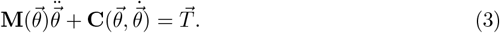

The moment of inertia tensor, **M**, and the Coriolis and Centripetal moments, **C**, are listed in [18]. Active torques, 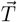, are generated by four monoarticular (musculus pectoralis and deltoid at the shoulder, biceps long head and triceps lateral at the elbow), and two biarticular muscles (biceps short head and the triceps long head). Muscle origins and insertions are taken from [19].

#### Force generation

Muscle force generation depends on muscle length, which is determined by the geometry of the arm and the current joint angles. We approximate this dependence linearly,

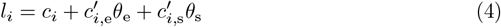

(*i* = 1, …, 6). The parameters, *c*_*i*_ (resting length) and 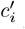 (moment arm) are taken from [19], and [20], and are listed in Appendix A. We neglect the second order terms of those sources, a reasonable approximation within the studied range of motion (see Figure 1b/c in [21]).

Given an activation signal, *A*_*i*_, for each muscle, *i*, force generation is specified in three steps. An exponential characteristic

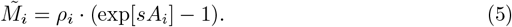

is weighted by the factor, *ρ*_*i*_, reflecting each muscle’s strength that is assumed to vary with the physiological cross-sectional area (PCSA) [19] (values listed in Appendix A). The factor *s* is constant across all muscles and based on empirical estimates from the cat gastrocnemius muscle [22]. In a second step, calcium kinetics is modeled as a critically damped second order low pass filter [11]

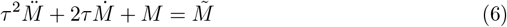

with a single parameter, *τ*, listed in Table 5. The third step takes into consideration the dependence of muscle force, *F*_*i*_, on the muscle contraction rate [23] and passive muscle elasticity,

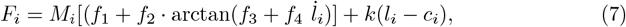

where *c*_*i*_ is the passive resting length of the muscle and the factors *f*_1_ to *f*_4_ and *k* were adjusted to reproduce the results of [11] (values listed in Appendix A).

The active joint torques, 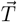, are obtained from the muscle forces, *F*_*i*_, taking the moment arms into account:

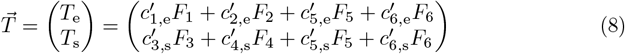

#### Reflex model

The activation, *A*_*i*_, of each muscle is assumed to reflect the Descending activation, *u*_desc,*i*_ acts as the threshold of a reflex loop that controls muscle activation, *A*_*i*_, for each muscle, *i* [11]:

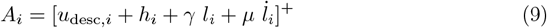

where [·]^+^ is a semi-linear threshold function. The parameters *γ* and *μ* reflect the sensitivity of muscle spindles to muscle length and its rate of change. The resting level of the *α*-motorneuron in the absence of descending activation is *h*_*i*_ *<* 0. Note that at rest (*i*_*i*_ = 0), descending activation (more precisely, − (*u*_desc,i_ + *h*)*/γ*) can be construed as a threshold length of the muscle as is done in the *λ*-model of posture [16]. When descending activation varies during movement generation, this interpretation is not straightforward as the rate of change of muscle length matters, the gains *μ* and *γ* matter, and it is not guaranteed that such a threshold length lies within the physiological range of the muscle. To side-step such issues, we avoid this interpretation.

The sensory signals, *l*_*i*_ and *i*_*i*_, are typically assumed to be delayed by about 20 ms over the physical state of the muscle. We neglected this delay for the purpose of the optimal control computation. Once the optimal descending activation had been determined, we verified that generating movements with these descending activation patterns from a model that included delays led to very similar movement trajectories.

#### Estimating descending activation by optimal control

To estimate a plausible pattern of descending activation, we minimize the change of descending activation required to bring the limb from an initial to a terminal postural state within a given movement time. Mathematically, we determine the time course of the vector, 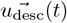, by minimizing the functional

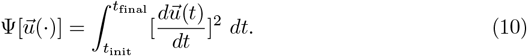

under the boundary conditions 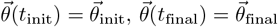 where *t*_final_ *t*_init_ = *MT* is the desired movement time. To guarantee the smoothness of the modeled movement, we also require that initial and terminal velocity and acceleration are zero:

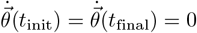 and 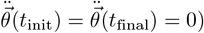 The minimization also imposes upper and lower bounds for joint angles and descending commands, and thus for muscle forces, in order to stay within biomechanical and physiological limits.

The optimization problem was solved numerically using the control vector parameterization approach [24, 25] as described in the Appendix C.

Because co-contraction of opposing muscles does not change the kinematics of movement we needed to specify levels of co-contraction. That happened by setting both the initial and the terminal level of co-contraction such that each muscle generated forces of 50 *N* (see Appendix B for more explanation).

#### Simulations

We estimated descending activation patterns by minimization for a total of 24 different movements. These movements included the eight movements probed in the experiment (Figure 2) at the two movement times imposed in the experiment (fast: 0.4 s; slow: 0.8 s). To study the transition from slow to fast movements, we also estimated descending activation patterns for an intermediate movement time (0.6 s). The movements generated by the model based on the estimated descending activation patterns were then systematically compared to the empirical movement data.

#### Estimating the virtual attractor trajectory

In the model, it is possible to directly compute the virtual attractor trajectories that are accessible empirically only through sophisticated instrumentation [17]. At any moment in time, the virtual attractor state is the set of joint angles and associated muscle lengths, at which the active torques contributed by all muscles sum to zero. In the model, we compute this state at each point in time given a current descending activation vector, 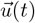, and current rates of change of all muscle lengths, 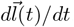.

Virtual attractor trajectories were obtained in joint space, but then transformed into hand position space using the kinematic model.

## Results

### Comparing model and experiment

#### Movement time

The movement times of the eight different movements performed at two instructed durations are listed in Table 1. Mean movement time across all movements and participants was 0.779 s in the slow condition and 0.445 s in the fast condition with standard deviations of 0.032 s and 0.022 s, respectively.

**Table 1.** Movement time (*T*) and its standard deviation (*T*_*SD*_) across participants for each of the eight movements in the two movement conditions, slow vs. fast.

| Movement | 1 | 2 | 3 | 4 | 5 | 6 | 7 | 8 |
| --- | --- | --- | --- | --- | --- | --- | --- | --- |
| slow |  |  |  |  |  |  |  |  |
| $T$ [sec] | 0.798 | 0.791 | 0.809 | 0.755 | 0.717 | 0.694 | 0.809 | 0.855 |
| $T_{SD}$ [sec] | 0.025 | 0.018 | 0.030 | 0.023 | 0.018 | 0.019 | 0.019 | 0.019 |
| fast |  |  |  |  |  |  |  |  |
| $T$ [sec] | 0.455 | 0.448 | 0.460 | 0.443 | 0.400 | 0.495 | 0.482 | 0.479 |
| $T_{SD}$ [sec] | 0.025 | 0.018 | 0.030 | 0.023 | 0.018 | 0.019 | 0.019 | 0.019 |

#### Kinematics

There is nothing new or surprising about the kinematics of the experimental data that come from a standard movement task. Hand paths (Figure 3) are relatively straight, hand trajectories (Figure 4) smooth and hand velocity profiles (Figure 5) bell-shaped. Joint trajectories and velocities (not shown) are likewise smooth as reported in the past [26].

**Fig 3.**
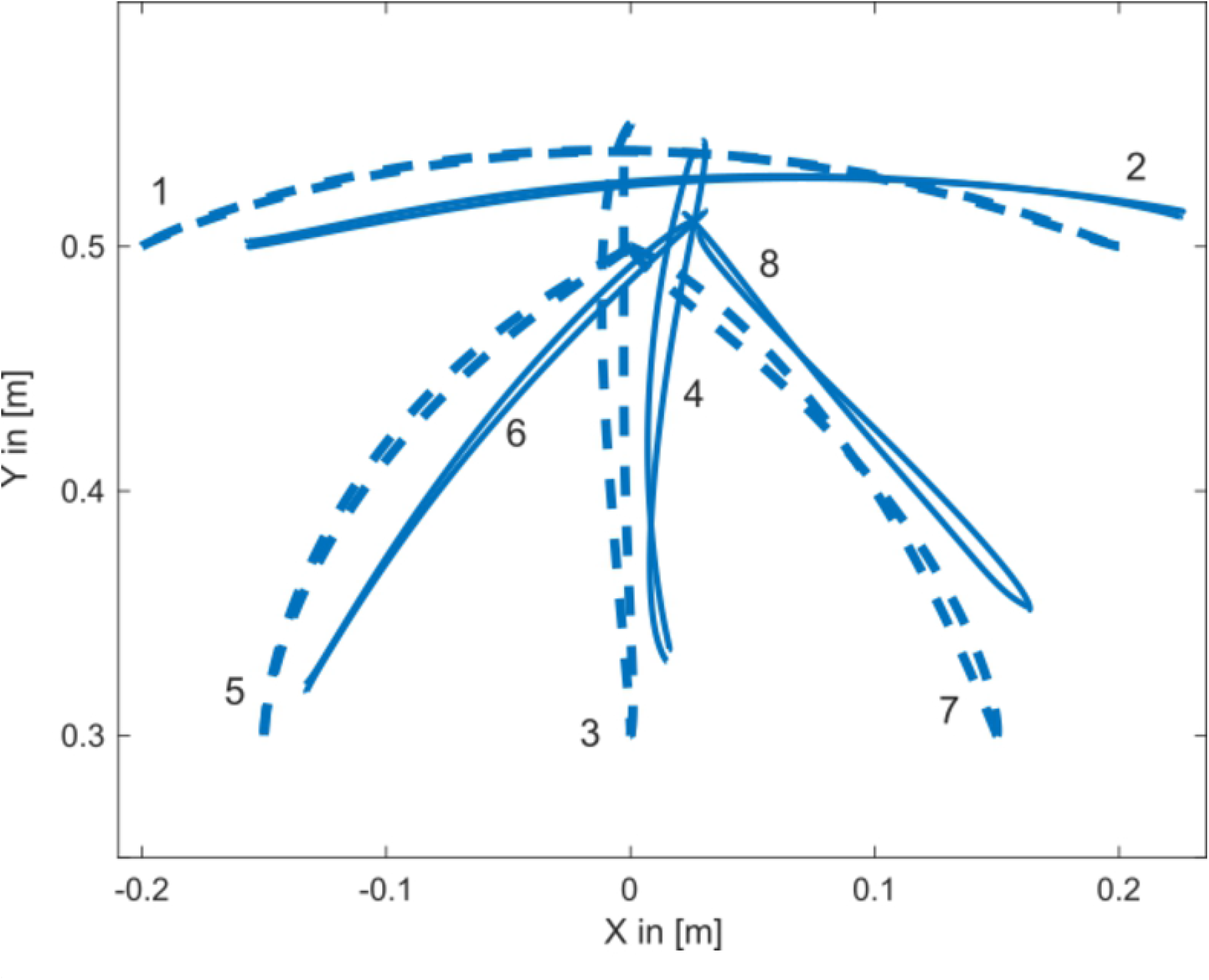
Hand paths from experiment (solid line, means across trials and participants) and from the model (dashed line) are shown for the slow condition (800 ms). The movement conditions are labeled as in Figure 2).

**Fig 4.**
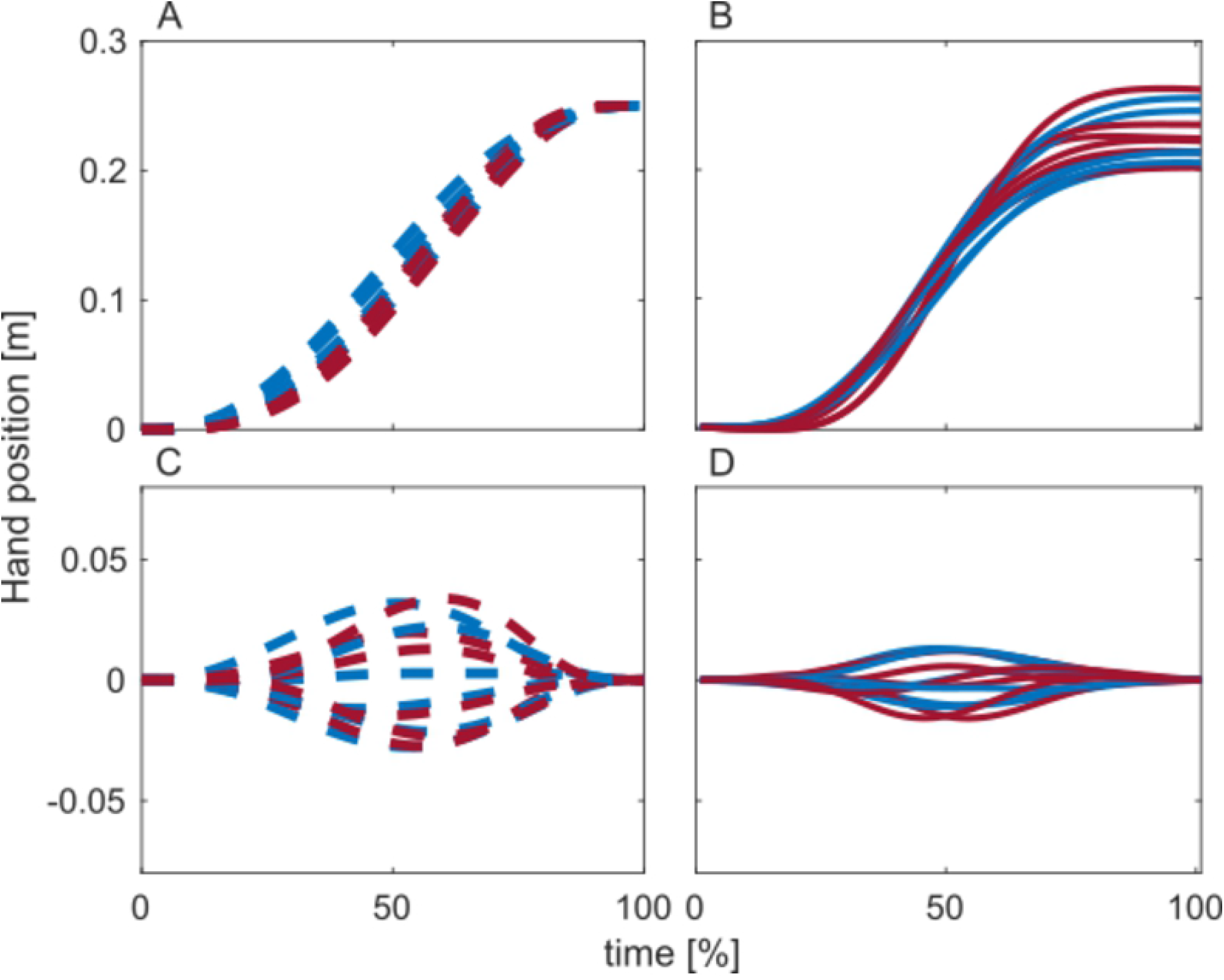
The hand trajectories of the six small amplitude movement conditions (3-8) for fast (red) and slow (blue) movements from the model (left column) and from experiment (right column). The top row shows the hand’s position along the line linking initial to target position, the bottom row shows the hand’s position orthogonal to that line. Experimental data (right column) are shown as means across participants and trials.

**Fig 5.**
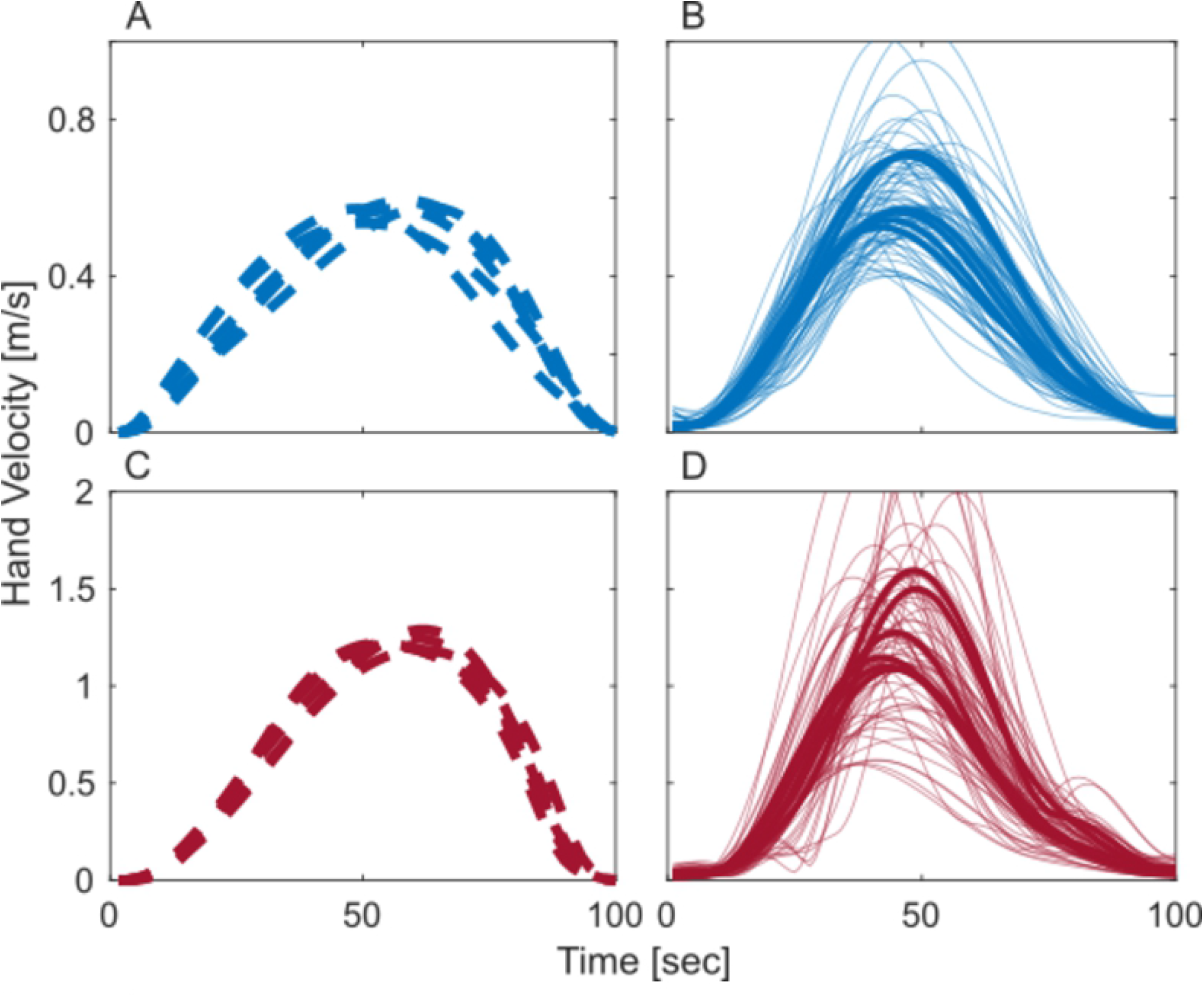
The hand velocity (length of the velocity vector in hand-space) of the six small amplitude movement conditions (3-8) from the model (left column) and from experiment (right column). The top row shows slow, the bottom row fast movements. Experimental data (right column) are shown as means across participants and trials (fat lines) and as means across trials for all participants and movement conditions (thin lines).

Hand paths from experiment match those from the model qualitatively (Figure 3). For all movement conditions, the small deviation from a straight path was consistent in direction and amount of curvature between model and experiment. Because participants have different segment lengths and were not, therefore, identically positioned relative to the targets, assembling mean paths required alignment across participants. This was done by centering all hand paths in the shoulder position. As a result, experimental paths are shifted relative to the simulated paths.

The two-dimensional hand trajectories through space are shown in Figure 4 for the the two matching movement times conditions of experiment and model. Normalizing time for the two conditions leads to invariant hand trajectories in both experiment and model, which is remarkable given the different mechanical conditions at the two speeds. Model and experiment fit qualitatively. Note that larger differences in the experimental trajectories at the end of the movement reflect primarily shifted initial postures of participants with different segment lengths.

Similarly, the model matched the experimentally observed hand velocity profiles qualitatively 5. Again, normalizing time makes these profiles approximately invariant across movement speeds. A small, but systematic difference between the experimental and simulated velocity profiles is the form of time asymmetry, especially at higher speeds. In experiment, hand velocity profiles rise faster and fall off slower than required by symmetry. In the model, this asymmetry is reversed.

#### Kinetics

The time courses of joint torques shown in Figure 6 provide a highly sensitive measure of the fit between model and experiment. The overall temporal profiles of torques are similar across movements, with profiles of reverse movements (2, 4, 6, 8) approximately mirrored relative to the profiles of forward movements (1, 2, 5, 7). Torques in the fast condition are essentially scaled in amplitude (in normalized time!) over torques in the slow condition.

**Fig 6.**
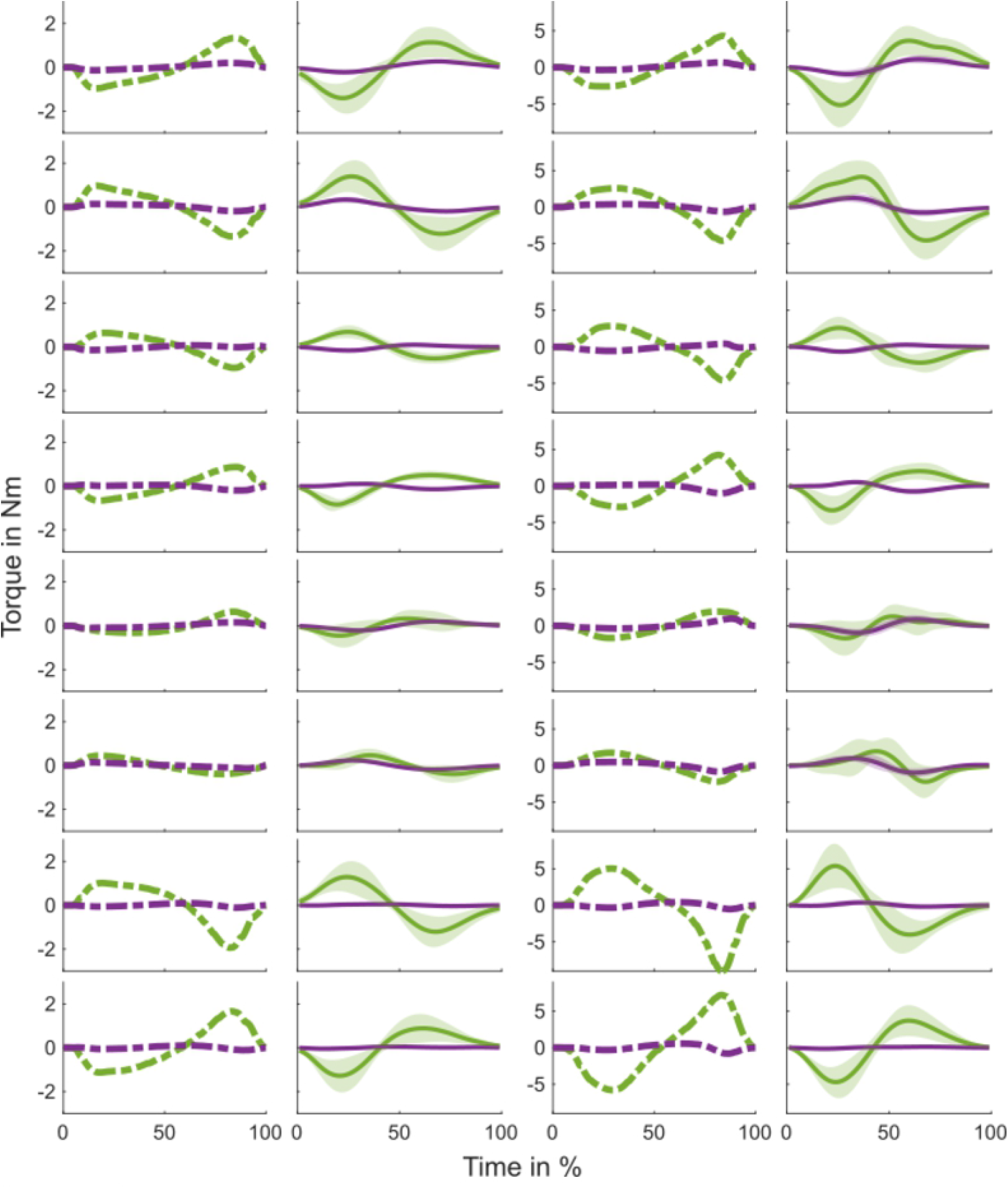
Simulated and experimental active joint torques across normalized time for all eight movements (rows) for the shoulder (green) and elbow (lilac). The first pair columns show the slow condition, the second pair of columns shows the fast condition. In each pair, the simulation comes first, the experiment second. The shaded areas in the plots of the experimental data shows the standard deviation.

The overall shape and scale of the torques from the model are similar to the experimental torques, Figure 6. However, the model differs again on asymmetry. While experimental torques profiles are close to symmetric or have a shorter first peak phase–longer second peak phase, the model torques profiles tend to have have the reserve asymmetry.

### Examining the estimated solutions

#### Time courses of descending activation

The time courses of the descending activation estimated by optimization are shown in Figures 7 for the three flexor muscles and in 8 for the three extensor muscles across all eight movements and three speed conditions of the model. The temporal profiles vary, but four patterns can be recognized, described here in order of increasing complexity: (1) In a small number of cases, descending activation is essentially constant in time and invariant across speed (e.g., for elbow flexors and extensors in movements 1 and 2; the biarticular flexors in movements 7 and 8). This is associated with little movement for the associated joint (movements 1 and 2) or counter-directional movements of the two joints for biarticulatory muscles (movements 7 and 8). (2) In another small number of cases, a ramp change extends across the entire movement, largely invariant under speed (e.g., shoulder extensor movement 2; elbow extensor movement 4; biarticular flexor movement 1 and 5). This is typically associated with a redundant muscle co-varying in a more complex form (e.g, the biarticular flexor covaries with the shoulder flexor in movement 1 but has a complex and speed dependent shape). (3) A common pattern is a ramp-like change that reaches its terminal level at about 70% of movement time and then stays constant, often seen at low speeds or across all speeds (e.g., shoulder and biarticulatory flexors and extensors in movement 1; elbow and biarticulatoy flexors in movement 4; and many others). (4) At high speeds, descending activation profiles often become non-monotonic, reaching a peak at around 40 to 60% of movement time and then reversing toward the final value (e.g., biarticulatory extensor in movement 2; all extensors in movement 3; shoulder flexor in movement 1 and 4; and many more). This resembles the “N-shape” reported earlier in the literature [27], although descending activation never returns across the terminal level. This may reflect muscles being more strongly driven by descending activation while accelerating then while breaking (see below).

**Fig 7.**
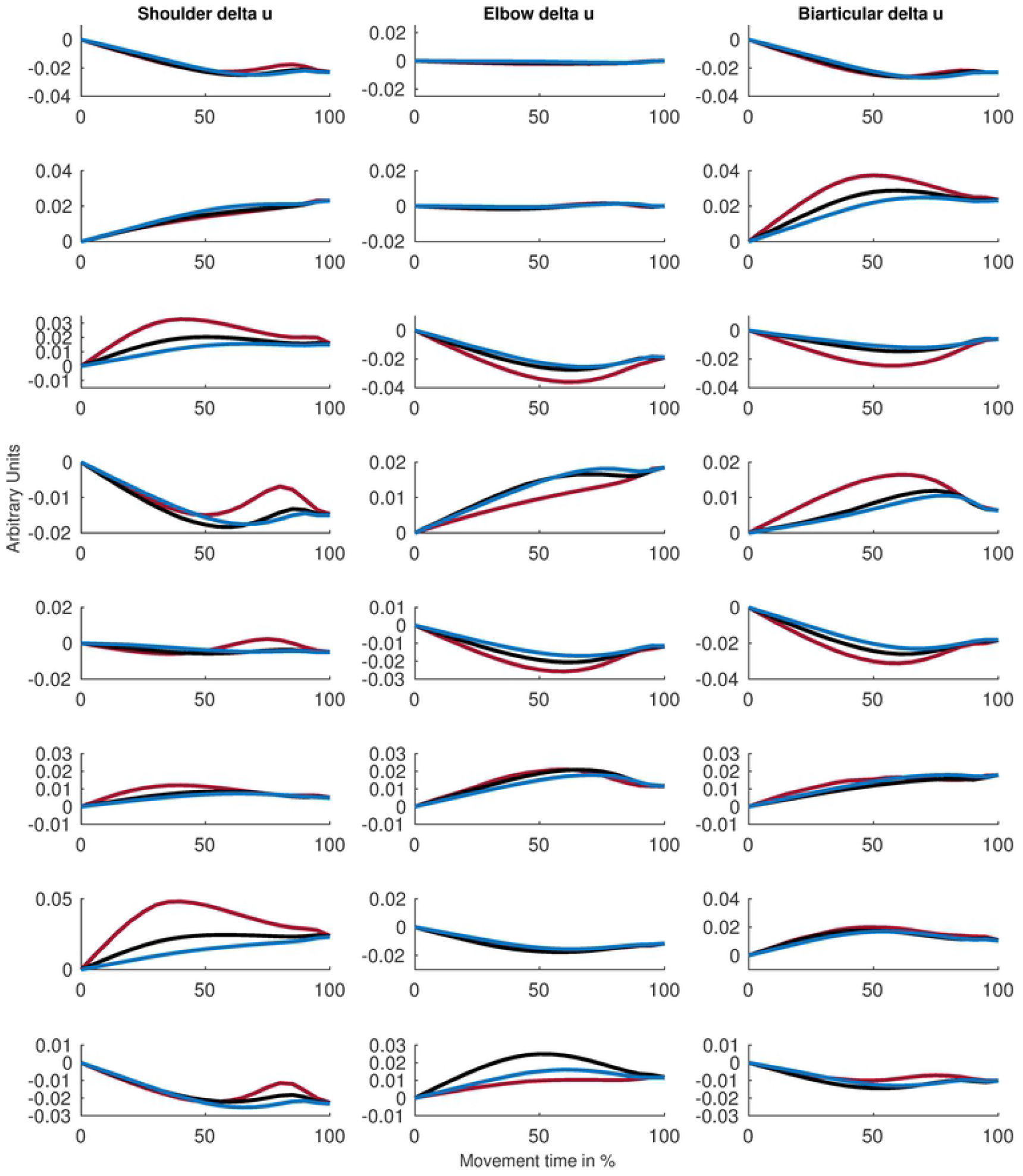
Time courses obtained in the model for the descending activations of the three flexor muscles for all eight movements (rows) and three speeds (blue: slow, black: mid; red: fast) are plotted in normalized time. The three columns show, from left to right, the musculus pectoralis (shoulder), the biceps short head (elbow) and the biceps long head (biarticular).

### The virtual attractor trajectory

The virtual attractor trajectory computed from the model in hand-space is shown in Figure 9 together with the predicted hand trajectories. The following five observations can be made. (1) As shown earlier (Figure 5, the model’s hand trajectories are invariant across speed and reflect a relatively straight path with only a small component orthogonal to movement direction (bottom row). (2) For slow movements (right column), the virtual attractor trajectory is largely aligned with the hand’s trajectory. Along movement direction (top), the virtual trajectory slightly leads during acceleration and falls back during deceleration. Orthogonal to movement direction, the virtual trajectory is constant and has little temporal structure. (3) For fast movements (left column), the virtual attractor trajectory along movement direction (top) is clearly N-shaped, strongly ahead of the hand’s trajectory early and then strongly fall back behind the hand’s trajectory late. Orthogonal to movement direction (bottom), the virtual attractor trajectory is temporally structured with larger amplitude than the hand’s trajectory. (4) The virtual attractor trajectory is, therefore, not invariant across movement speed while than hand’s trajectory is. (5) At a given speed, the virtual attractories are quite invariant across the different movements, more so than the temporal profiles of the descending activation patterns (Fig. 7, 8).

**Fig 8.**
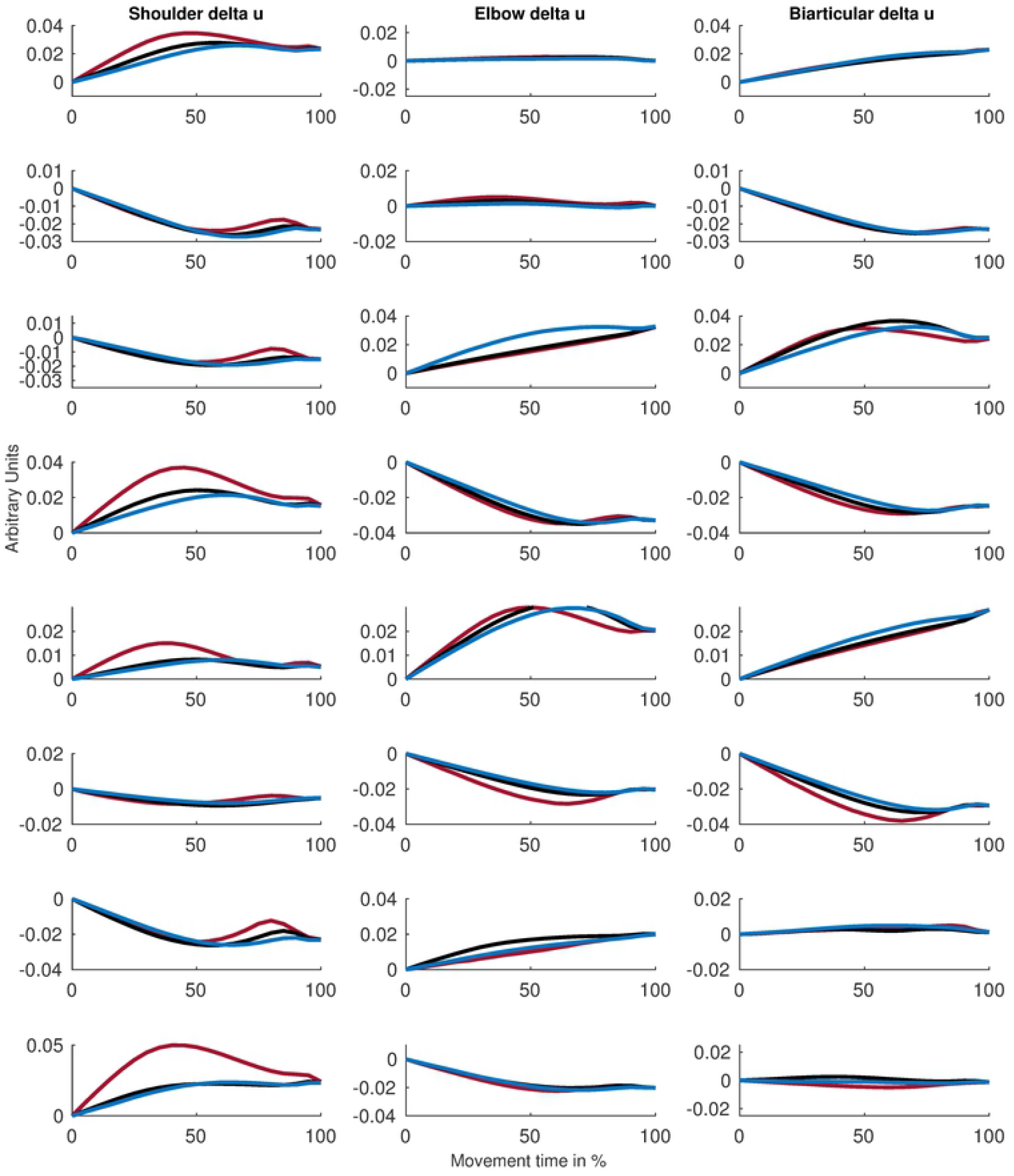
Time courses obtained in the model for the descending activations of the three extensor muscles, from left to right: deltoid (shoulder), the triceps lateral head (elbow) and the triceps long head (biarticular). All other conventions as in Figure 7.

**Fig 9.**
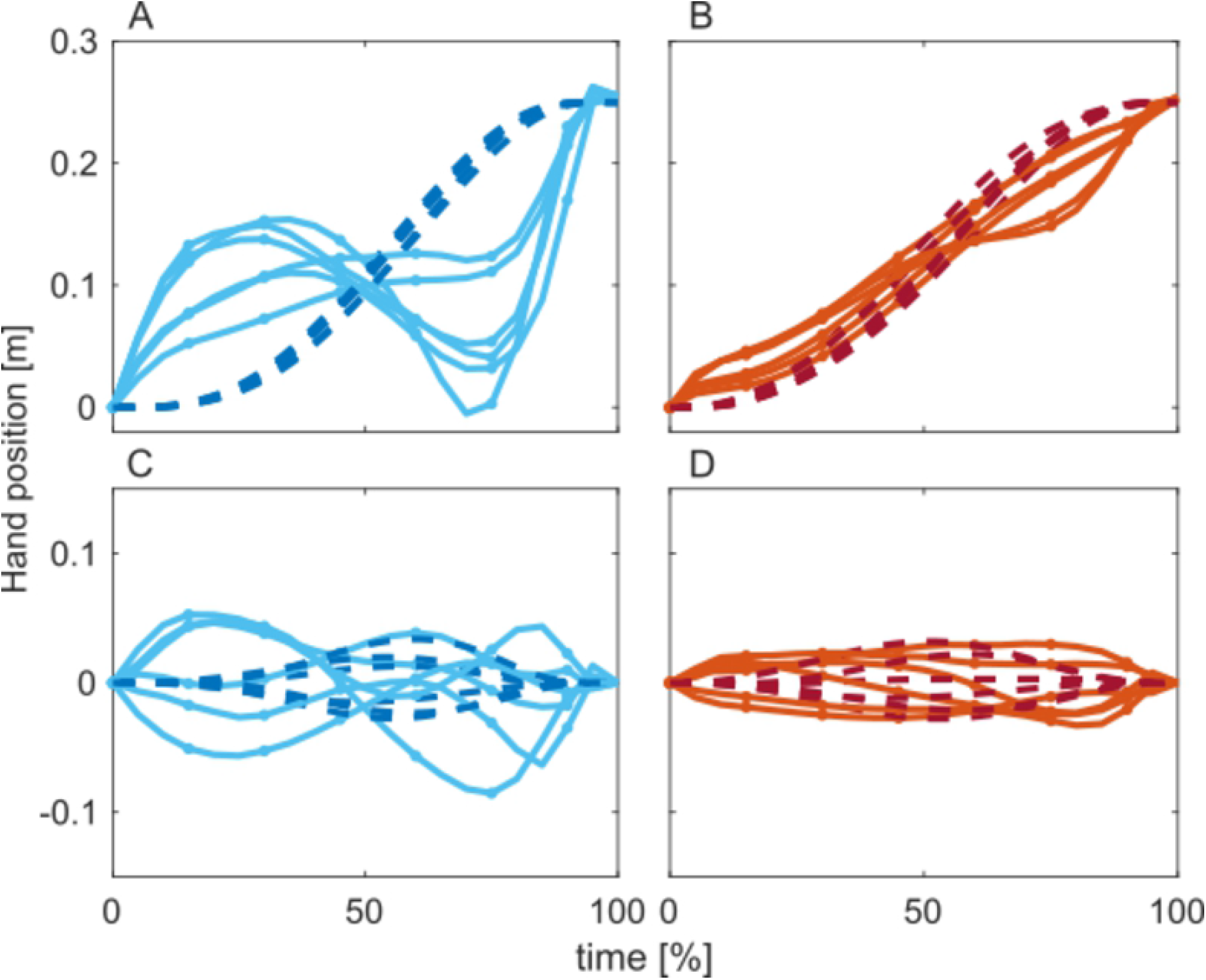
The hand-trajectories obtained from the model (dashed, dark) for movements 3-8 are plotted in normalized time along the line connecting the initial to the target posture (top) and orthogonal to that line (bottom). Overlaid are the corresponding attractor trajectories (solid, light). The left column depicts the fast, the right column the slow condition.

## Discussion

### How well does the model capture human movement data?

We compared kinematics and kinetics of the movements predicted by the minimal descending activation patterns to human movements observed experimentally at two different movement times. The movements generated by the model qualitatively match observed hand movement paths, trajectories, and velocity profiles quite well. This includes catching the invariance of the hand’s kinematics across a doubling of movement speed. The model’s prediction also match observed joint torque profiles and their scaling with different movement times.

A limitation of the match to experiment lies in the temporal asymmetry of different state variables. Human movement trajectories are slightly asymmetrical, especially at higher speeds, with a shorter and steeper acceleration phase compared to the longer and slower deceleration phase. This pattern is inverted in the model.

Generally, the model of the stretch reflex, Equation 9, implies that when a muscle lengthens, the same amount of descending activation generates more muscle activation than when a muscle shortens. Because acceleration is caused by muscles that shorten and breaking is caused by muscles that lengthen, this implies that more change of descending activation is needed to accelerate than to break a movement. Minimizing descending activation change thus leads to slower/lower acceleration than deceleration. Muscle physiology would tend to compensate for this effect as contracting muscles produce more force than lengthening muscles, reducing the amount of descending activation needed to bring about acceleration. Hill type muscle models capture this physiological effect [20, 28], and would thus reduce or potentially reverse this discrepancy. In contrast, the simplified model of muscle force generation that we used [11], amplifies the discrepancy because in this model, the same level of muscle activation generates more force for lengthening than for shortening muscles. We chose this simplified model to enable comparison to that earlier work (see below). The simplified model also sidesteps the additional redundancy implied in Hill-type models by the non-unique mapping between the length of the contractile element and the joint angle.

### What does the cost function mean?

The capacity of the model to predict the observed kinematics and its invariance across rate and workspace is remarkable given that no physical or physiological factor enters into the cost function. This is in contrast to other approaches, such as minimal torque change [29] and minimum effort [13]. The latter model in effect minimizes muscle activation. The asymmetry of the velocity profiles is not consistently captured by that model either (see their Figure 4b, which is not directly compared to experiment, however).

Thus, merely minimizing the temporal complexity of the descending activation pattern captures some of the regularity of human movement kinematics. This is consistent with the general theme that the peripheral reflex circuitry and muscle dynamics simplify motor control [10, 30].

This theme was at the center of the modeling study from which we adopted the reflex and simplified muscle model [11]. That study showed that simple “ramp” descending activation patterns (framed by those others as virtual trajectories) are sufficient to generate realistic movement patterns and stiffness profiles. Our approach directly speaks to that observation. In fact, the optimal control problem as framed here leads to such ramp-like solutions in the limit case of a trivial plant. Interpreting the descending activation as threshold muscle length, *λ* [16], the trivial plant is realized when muscle lengths simply track threshold lengths, *l*(*t*) = *λ*(*t*). In that case, the optimal control problem can be easily solved analytically and leads to a linear ramp in *λ*(*t*) that connects the initial to the final value for each muscle. Since we initialized optimizations with exactly this linear ramp, the resulting descending activation patterns results may indeed be interpreted as the solutions closest to the linear ramps while compatible with the imposed movement target and time. These solutions deviate from the linear ramp because the real plant deviates from the trivial one due to biomechanics, muscle dynamics, and reflex delays.

The ramp-like profiles we found for slow movements approximates the limit case of the trivial plant, although the ramp ends before the movement ends. This is consistent with experimental estimates of the time when the descending signal reaches its terminal level [31]. Those experiments actually involved fast movements, and did not estimate the temporal profile of the descending activation pattern. Our estimates for fast movements identified temporally more complex, non-monotonic descending activation patterns. Earlier work hypothesizing ramp-like profiles for the same neuro-muscular model used here [11] did not explore movement speeds and portions of work space at which we found these non-monotonic estimates. Thus, the modeling work reported here clarifies earlier debates about the complexity of descending signals required to bring about movement.We do not, however, seriously propose that the nervous system minimizes the temporal complexity of descending activation patterns. The optimal control approach was used primarily as a tool to provide reasonable estimates of the descending activation patterns, which we may then analyze for their regularities and properties.

### The temporal structure of descending activation patterns

We discovered a range of different time structures of the minimally changing descending activation patterns (Fig. 7, 8). Slow movements tended to show constant-rate ramp-like profiles that reached the final level at about 60 to 70 percent of movement time. Faster movements tended to show activation patterns that were not monotonic, overshooting and then returning to the final level. These patterns may reflect the anticipation of interaction torques and delays in building the required muscle force levels, which increase in size and relevance of increasing movement speeds. In fact, the descending activation patterns of muscle pairs that contribute significantly to acceleration and deceleration of the movement acquire this more strongly modulated temporal structure for faster movements, consistent with the need to overcome inertial and interaction torques (Fig. 7 and 8). Qualitatively, these descending activation patterns resemble the “N-shapes” postulated in [27] (although that earlier work entailed single joints and an inadequate linear model as argued in [32]). Descending activation patterns of muscle pairs that contribute weakly to the acceleration and deceleration of motion tend to be strongly modulated in time.

The contribution of biarticular muscles that span the shoulder and elbow joints depends on the joint configuration and the direction of a movement. The descending activation patterns reflect these dependencies. In general, co-changes of elbow and shoulder were supported by descending activations to the biarticular muscles that matched the temporal profile of the activation descending to the monoarticular muscles (movements 5 and 6 in Figure 2, 5th and 6th row in Figures 7 and 8). In movements involving elbow and shoulder movements in opposite directions, the role of the biarticular muscles varies with the extent of either change. When activation descending to the monoarticular muscles have different amplitudes, then activation descending to the biarticular muscles share the temporal profile with the activation descending to the monoarticular muscles of larger amplitude (movements 1, 2, 3, 4). When the activations descending to the monoarticular muscles are about equal in size, then the activations descending to the biarticular muscles are approximately constant in time and do not contribute significantly to the movement (movements 7 and 8).

Co-contraction was specified for the initial and identically for the final state as a boundary condition for the optimization (see Appendix B). During the movement, co-contraction emerged from the optimization. By varying the imposed initial/final level of co-contraction we found that these levels had little influence on the descending activation patterns generated by the optimization and on the predicted movement.

The observed difference between temporal profiles of descending activation patterns of slow versus fast movements refutes the idea that movement generation can be explained by a single movement template or primitive, that would be scaled to the desired movement time. Figures 7 and 8) already use normalized time so a simple scaling would imply identical temporal structure for different movement speeds. When descending activation patterns are specifically shaped to anticipate velocity dependent torques and muscle properties, then the neural processes generating those temporal profiles could be considered to reflect an “internal model” as postulated in computational theories of motor control [6, 33].

### The virtual attractor trajectory

The time course of the joint angles and associated muscle lengths which all active torques generated by the muscles are zero is called the virtual attractor trajectory by [17]. Unlike in experiment, it is easy to determine these virtual attractor trajectories in the model. We found these to be astonishingly uniform across the different movements when plotted in end-effector space in coordinate systems that are aligned with each movement (Figure 9). Along the movement, the virtual attractor trajectories of slow movements are linear ramps that reach all the way to the end-point of the movement. Within increasing speed, the virtual attractor trajectories become increasingly “N-shaped”, initially leading the hand’s position to accelerate and then falling behind the hand’s position to decelerate. Orthogonal to the movement direction, the virtual attractor trajectories are approximately constant in time, with a small modulation at higher movement speeds that may reflect compensation of interaction torques.

It is important to note that all virtual attractor trajectories lie within the reachable work space. Thus, at every moment in time during a movement, a realisable muscle length exists for all muscles at which descending activation is compensated for by activation from the stretch reflex (neglecting co-contraction). If the hand was to exactly trace the virtual trajectory (as brought about in the experiments of [17] by applying exactly the right external torques), activation that descends would be exactly equal to the activation that the reflex loop contributes (again, except for co-contraction). In a sense, the distance between the real and the virtual hand trajectory reflects the difference between these two contributions. The observed patterns suggest that the contribution of the reflex to movement generation is of the same order of magnitude as the contribution of descending activation itself, especially for slow movements.

It may be tempting to think of the invariant shape of the virtual trajectory as a simple control strategy. Note, however, that the descending activation patterns that the brain’s neural networks must generate are less invariant than the virtual trajectories that reflect the contributions of biomechanics, muscle dynamics, and reflex loops. The invariance of the virtual trajectory thus essentially is the dynamic reflection of the invariance of human movement kinematics across work space and speed.

### Limitations

Clearly, this model provides only a first rough approach to using models of the periphery to estimate descending activation patterns. Using more realistic and detailed muscle models and including more complete accounts of spinal reflexes is a future challenge [10].

Muscle redundancy was also only addressed summarily, but minimizing the change of activation to redundant muscles. A more thorough-going analysis of muscle redundancy may require new experimental approaches to enable estimation of the descending activation of individual muscles. Similarly, kinematic redundancy [34] adds an important next level of complexity that would need to be addressed by the approach we sketched here.

For computational reasons, we neglected delays in the reflex loop. To assess the error induced, we input the estimated descending activation patterns into a model that included typical sensory delays (as quantified in [11]). We found that the resultant kinematics differ little from the kinematics of the model without delay. This is likely due to the fact that in the most dynamic portions of the movement, early acceleration and late deceleration, joints and muscle lengths vary slowly.

### Conclusion

Taking biomechanics, muscle dynamics, and the stretch reflex into account, we determined the descending activation patterns that change minimally to bring about targeted movements at different speeds. These patterns predicted observed human movement trajectories quite well. The temporal structure of these descending activation patterns changes from monotonic for slow movements to increasingly complex and non-monotonic for faster movements. Thus, movements at different speeds cannot be explained by a simple re-scaling of descending activation templates. The time structure of descending activation reflects the variation with speed of interaction torques and muscle dynamics, consistent with the role of the internal models postulated in computation theories of motor control. Attractor trajectories, on which active muscle torques are zero, reflect the balance of contributions from descending activation and the stretch reflex. Attractor trajectories were relatively invariant across movements when analyzed in terms of the hand’s trajectory along and orthogonal to movement direction. They lie within reachable work space, and reflect similar orders of magnitudes of descending and reflex-generated activation during movement.

## Acknowledgements

We gratefully acknowledge helpful discussion with Dr. Lei Zhang.

## Author contributions

Conceptualization: RR, JSJ, CH, GS

Data Curation: RR, CH

Formal Analysis: RR, CH

Investigation: RR, CH

Methodology: RR, CH, JSJ, GS

Project Administration: GS

Resources: GS

Software: RR, CH, JSJ

Supervision: GS, JSJ

Validation: RR, CH

Visualization: RR, CH

Writing – Original Draft Preparation: RR, CH, GS

Writing – Review & Editing: RR, CH, GS

## A Model parameters

Values of all parameters of the model are listed in the following Tables 2 to 5.

**Table 2.** Moment arms and resting lengths of the muscles are taken from [20]. Muscle lengths are taken from [19].

| muscle | $c'_{i,s}$ in [m] | $c'_{i,e}$ in [m] | $c_i$ in [m] |
| --- | --- | --- | --- |
| elbow flexor ($l_1$) | 0.000 | -0.014 | 0.287 |
| elbow extensor ($l_2$) | 0.000 | 0.025 | 0.246 |
| shoulder flexor ($l_3$) | -0.030 | 0.000 | 0.216 |
| shoulder extensor ($l_4$) | 0.030 | 0.000 | 0.191 |
| biarticular flexor ($l_5$) | -0.030 | -0.016 | 0.333 |
| biarticular extensor ($l_6$) | 0.030 | 0.030 | 0.312 |

**Table 3.** Values for the biomechanical parameters of the lower and the upper limb are taken from [19].

|  | length [m] | mass [kg] | inertia [Nm] |
| --- | --- | --- | --- |
| upper arm | 0.3348 | 2.1 | 0.0244 |
| lower arm | 0.2628 | 1.2 | 0.0076 |

**Table 4.** Values for the physiological cross section areas for the six modeled muscles are taken from [19].

| muscle | PCSA [cm<sup>2</sup>] |
| --- | --- |
| $\rho_1$ | 2.10 |
| $\rho_2$ | 4.52 |
| $\rho_3$ | 4.52 |
| $\rho_4$ | 3.87 |
| $\rho_5$ | 1.29 |
| $\rho_6$ | 3.87 |

**Table 5.** Values of parameters that specific to the neuro-muscular model were taken from [11] and adjusted such that the results of [11] could be reproduced. Parameters *f*_1_, *f*_2_, *f*_3_ *f*_4_ and *c* are fitting parameters for the active and viscous part of force generation. The passive stiffness is scaled with *k. τ* is the time constant of the differential equation for the muscle force generation.

| parameter | value |
| --- | --- |
| $f_1$, [-] | 0.82 |
| $f_2$, [-] | 0.57 |
| $f_3$, [-] | 0.43 |
| $f_4$, [s/m] | 58 |
| $\tau$, [s] | 0.015 |
| $k$, [-] | 17.35 |
| $s$, [1/m] | 112 |

## B Co-contraction

The amount of co-contraction, *C*, specifies how much all threshold lengths, 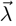, must be shifted such that the forces generated in equilibrium by individual muscles reach that given level without changing net joint torques. We separate the descending signals 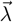 into a reciprocal part 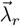 and a co-contraction part 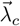 with 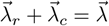. Furthermore, we consider the stationary state without external torques. Under those conditions, the reciprocal part of the reference command equals the actual muscle lengths, 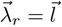, and does not contribute to the generation of forces. To determine the co-contraction component of the reference command, 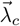, we compute the threshold lengths at which the force generated by each muscle is increased by an average of *C* (in Newton) by solving:

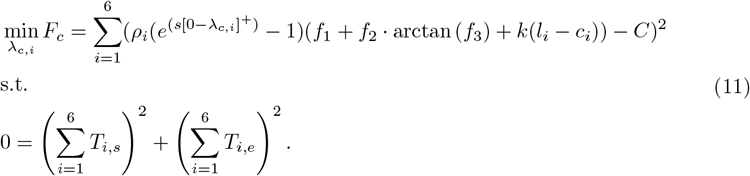

Here, *T*_i,e_ and *T*_i,s_ are the elbow and shoulder torques induced by the *i*-th muscle.

## C Control vector parameterization

We used the control vector parameterization approach [24] to optimize the time course of the reference commands 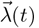. Each of the six continuous reference commands, *λ*_*i*_, was sampled at *N* = 20 moments in time, leading to a total number of 120 parameters to optimize. Optimization was performed with respect to the objective function defined in section using the iterative nonlinear programming solver `fmincon`. Initialization was chosen so that the reference commands sampled a linear ramp from the initial to the target position in the Cartesian plane. After each optimization step, cubic splines were used to interpolate the reference commands between the discrete sample points to provide time continuous input to the model.

In order to evaluate the objective function, we defined the 18-dimensional state trajectory 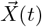, that completely describes the time dependent state of the arm during movement. It comprises the six ungraded muscle force profiles, 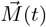, their rates of change, 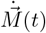, and the joint position, 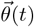, and velocity, 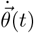, time courses during the movement.

The state trajectory, 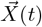, of a movement is obtained by solving the differential equations, 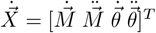 (obtained from equations 3 and 6), for each of the *N* time intervals of length *MT/N*. The final state of the (*i* − 1)th interval was used as the initial state for the *i*th interval. The initial state for the first integration interval is composed of the initial muscles states at a predefined level of co-contraction, 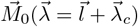 (see B), the initial joint positions 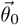, and zero rates of change (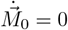 and 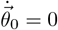). State trajectories were computed using the Matlab solver ode23tb.

